# fcfdr: an R package to leverage continuous and binary functional genomic data in GWAS

**DOI:** 10.1101/2021.10.21.465274

**Authors:** Anna Hutchinson, James Liley, Chris Wallace

## Abstract

**Summary:** GWAS discovery is limited in power to detect associations that exceed the stringent genome-wide significance threshold, but this limitation can be alleviated by leveraging relevant auxiliary data. Frameworks utilising the conditional false discovery rate (cFDR) can be used to leverage continuous auxiliary data (including GWAS and functional genomic data) with GWAS test statistics and have been shown to increase power for GWAS discovery whilst controlling the FDR. Here, we describe an extension to the cFDR framework for binary auxiliary data (such as whether SNPs reside in regions of the genome with specific activity states) and introduce an all-encompassing R package to implement the cFDR approach, `fcfdr`, demonstrating its utility in an application to type 1 diabetes.

**Availability and implementation:** The `fcfdr` R package is freely available at: https://github.com/annahutch/fcfdr. Scripts and data to reproduce the analysis in this paper are freely available at: https://annahutch.github.io/fcfdr/articles/t1d_app.html

## 1 Introduction

A stringent significance threshold is required to identify robust genetic associations in GWAS due to multiple testing constraints. Leveraging relevant auxiliary data has the potential to boost statistical power to exceed the significance threshold. The conditional FDR (cFDR) is a Bayesian FDR measure that additionally conditions on auxiliary data to call significant associations. The cFDR approach was originally developed to leverage GWAS *p*-values from related traits, thereby exploiting genetic pleiotropy to increase GWAS discovery ^1,2,3^, and has been shown to increase power for GWAS discovery whilst controlling the frequentist FDR ^11^.

Motivated by the enrichment of GWAS SNPs in particular functional genomic annotations ^14^, Flexible cFDR was developed to extend the usage of the cFDR approach to the accelerating field of functional genomics ^9^. However, at-present no cFDR methodology exists that permits binary auxiliary data, meaning that the approach cannot currently be used to leverage auxiliary data with a binary representation, such as whether SNPs are synonymous or non-synonymous or whether they reside in regions of the genome with specific activity states.

Here we present an extension to the cFDR approach that supports binary auxiliary data and we thus introduce a cFDR toolbox in the form of an R package (https://github.com/annahutch/fcfdr) that supports various auxiliary data types. We demonstrate the utility of our methods and software by iteratively leveraging three distinct types of relevant auxiliary data with GWAS *p*-values for type 1 diabetes (T1D) ^12^ to uncover new genetic associations.

## 2 The cFDR framework

Let *p*_1_, …, *p*_*m*_ ∈ (0, 1] be a set of *p*-values corresponding to the null hypotheses of no association between the SNPs and a trait of interest (denoted by *H*_0_). Let *q*_1_, …, *q*_*m*_ be auxiliary data values corresponding to the same *m* SNPs. Assume that *p* and *q* are realisations of random variables *P, Q* satisfying:

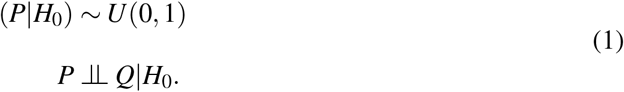

The cFDR is defined as the probability that a random SNP is null for the trait given that the observed *p*-values and auxiliary data values at that SNP are less than or equal to values *p* and *q* respectively ^1,2^. Bayes theorem and standard probability rules are used to derive:

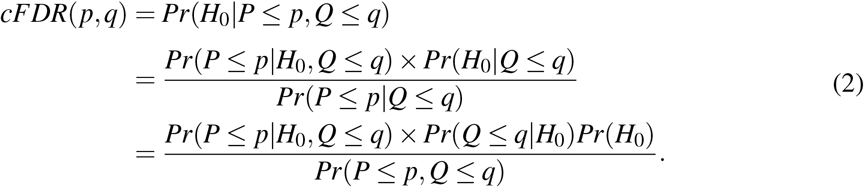

To construct a conservative estimator of the cFDR, approximate *Pr*(*P* ≤ *p*|*H*_0_, *Q* ≤ *q*) ≈ *p* (from property 1; note that if property 1 holds and *P* is correctly calibrated then this approximation is an 45 equality) and *Pr*(*H*_0_) ≈ 1 (since associations are rare in GWAS):

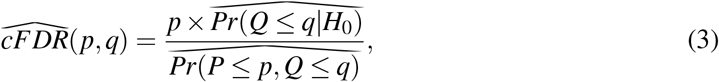

where ^ is used to denote that these are estimates under the assumption *H*_0_ ⫫ *Q*|*P*. The methods used to estimate the cumulative densities in equation (3) vary across approaches. In the original cFDR approach they are estimated using empirical cumulative density functions ^1,10,11^ whilst in Flexible cFDR they are estimated using kernel density estimation ^9^.

However, the 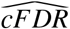 values do not directly control the FDR ^10^. Instead, a method proposed by Liley and Wallace ^11^ can be used to generate *v*-values, which are essentially the probability of a newly-sampled realisation (*p, q*) of *P, Q* attaining an as extreme or more extreme 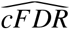 value than that observed, given *H*_0_. The *v*-values are therefore analogous to *p*-values and can be used in any conventional error-controlling multiple testing procedure that allows for slightly dependent *p*-values (e.g. the Benjamini-Hochberg procedure). The derivation of *v*-values also allows for the method to be applied iteratively to incorporate additional layers of auxiliary data.

Since binary auxiliary data can only take two values, we introduce an alternative methodology called “Binary cFDR” which is based on finding optimal rejection regions to derive *v*-values (Supplementary Methods). We show in a simulation-based analysis that applying Binary cFDR iteratively over informative auxiliary data increases power whilst controlling the frequentist FDR (Supplementary Results, Supplementary Fig. 2).

## 3 R package and T1D application

We present an R package that implements both Flexible cFDR and Binary cFDR, named `fcfdr` (https://github.com/annahutch/fcfdr), and demonstrate its utility in an application to T1D which is fully reproducible (https://annahutch.github.io/fcfdr/articles/t1d_app.html).

We used *p*-values from an Immunochip study of T1D ^12^ as our primary data set. In the first iteration we used Flexible cFDR to leverage Immunochip *p*-values for a genetically related trait, rheumatoid arthritis (RA) ^6^. In the second iteration we used Binary cFDR to leverage data measuring SNP overlap with regulatory factor binding sites ^5,8,7^ and in the third iteration we used Flexible cFDR to leverage average enhancer-associated H3K27ac fold change values derived from ChIP-seq experiments conducted in T1D-relevant cell types ^4^ (Supplementary Methods) (Fig. 1).

**Figure 1:**
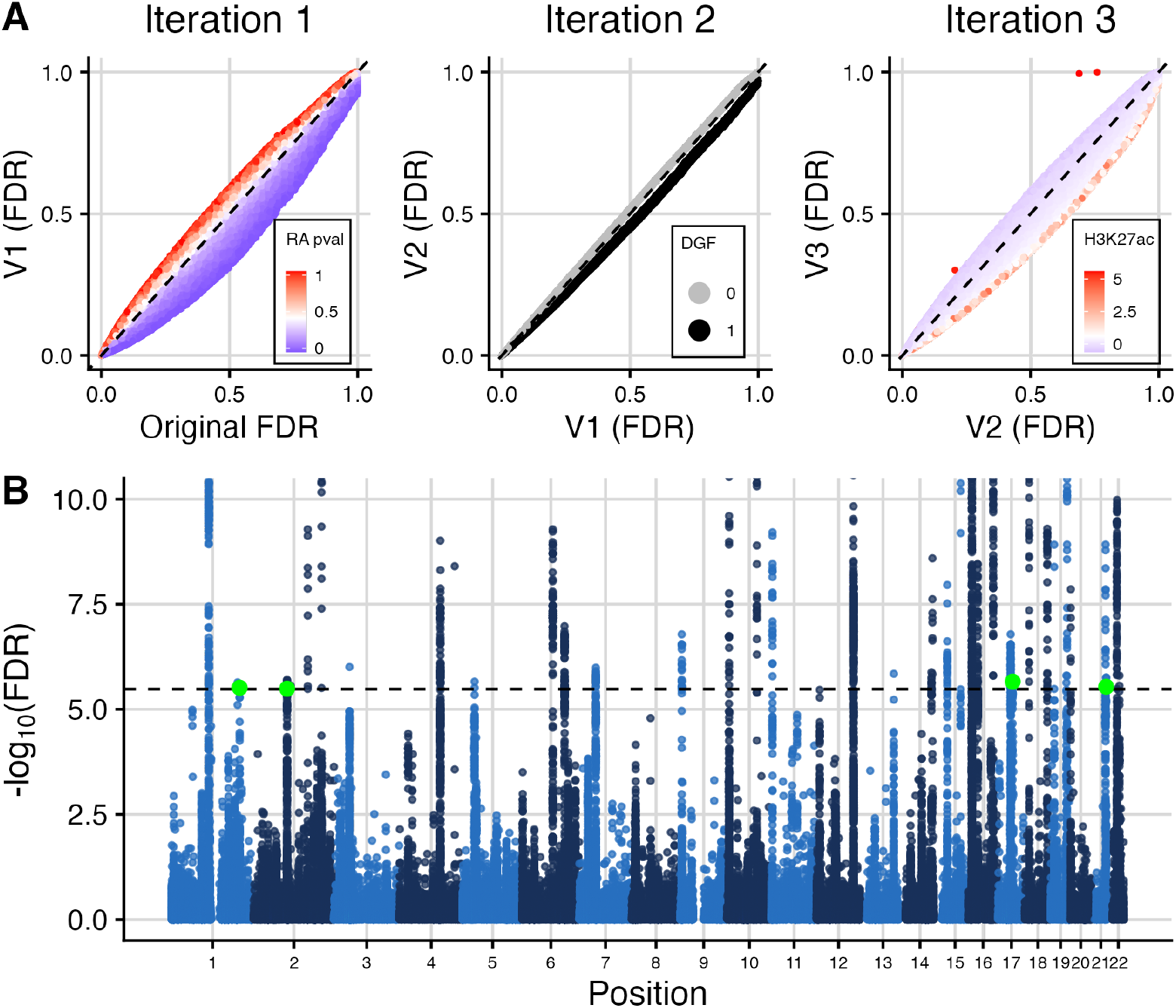
Summary of cFDR results for T1D application. (A) FDR values (derived from the Benjamini-Hochberg procedure) before and after each iteration of cFDR, coloured by the auxiliary data values. (B) Manhattan plot of (−*log*_10_) FDR values (*y*-axis truncated to aid visualisation). Green points indicate the four lead variants that were newly FDR significant after cFDR. Black dashed line at FDR significance threshold (*FDR* = 3.3*e* − 06).

Our implementation of cFDR identified 101 SNPs as newly genome-wide significant (*FDR* ≤ 3.3*e* − 06 which corresponds to *p* ≤ 5*e* − 08; Supplementary Methods). These SNPs had relatively small *p*-values for RA (median *p* = 0.007 compared with median *p* = 0.422 in full data set), were more likely to be found in regulatory factor binding sites (mean binary value was 0.406 compared to 0.234 in full data set) and had larger H3K27ac fold change values in T1D-relevant cell types (median fold change value was 1.44 compared with 0.576 in full data set). Similarly, 45 SNPs were identified as newly not significant (i.e. they were significant in the original GWAS data set but became not significant after applying cFDR). These SNPs had relatively high *p*-values for RA (median *p* = 0.620), were less likely to be found in regulatory factor binding sites (mean binary value was 0.044) and had smaller H3K27ac fold change values in T1D-relevant cell types (median fold change value was 0.431).

The original GWAS identified 38 significant genomic regions (based on our definition of genomic regions; Supplementary Methods). All of these were found to be significant in the cFDR analysis, which additionally identified 4 genomic regions that were newly significant (with lead variants: rs1052553, rs3024505, rs6518350 and rs13415583) (Fig. 1). Three of these SNPs had small *p*-values for RA (rs1052553: RA *p* = 0.007; rs6518350: RA *p* = 0.06161 and rs13415583: RA *p* = 1.913*e* − 06 whereas rs3024505 had RA *p* = 0.6008) and two of these SNPs had high H3K27ac fold change values (rs3024505 had 87.4th percentile and rs6518350 had 72.7th percentile of H3K27ac fold change values). Two of the lead variants overlapped regulatory factor binding sites (rs1052553 and rs3024505). When using a larger Immunochip study of T1D for validation (16, 159 T1D cases compared to 6, 670) ^13^, we found that three out of the four lead variants were present and that these had smaller *p*-values in the validation GWAS data set than the discovery GWAS data set: rs1052553 had *p* = 1.649*e* − 15, rs3024505 had *p* = 9.127*e* − 14, rs13415583 had *p* = 4.764*e* − 09 in the validation data set ^13^ compared to *p* = 8.156*e* − 08, *p* = 6.394*e* − 08 and *p* = 1.062*e* − 07 respectively in the discovery data set ^12^.

## 4 Conclusion

We have described a novel implementation of the cFDR approach that supports binary auxiliary data and have introduced an all-encompassing R package, `fcfdr`, that can be used to implement the cFDR approach for a wide variety of auxiliary data types. We have demonstrated the versatility of this tool in an application to T1D where we uncovered new genetic associations.

---

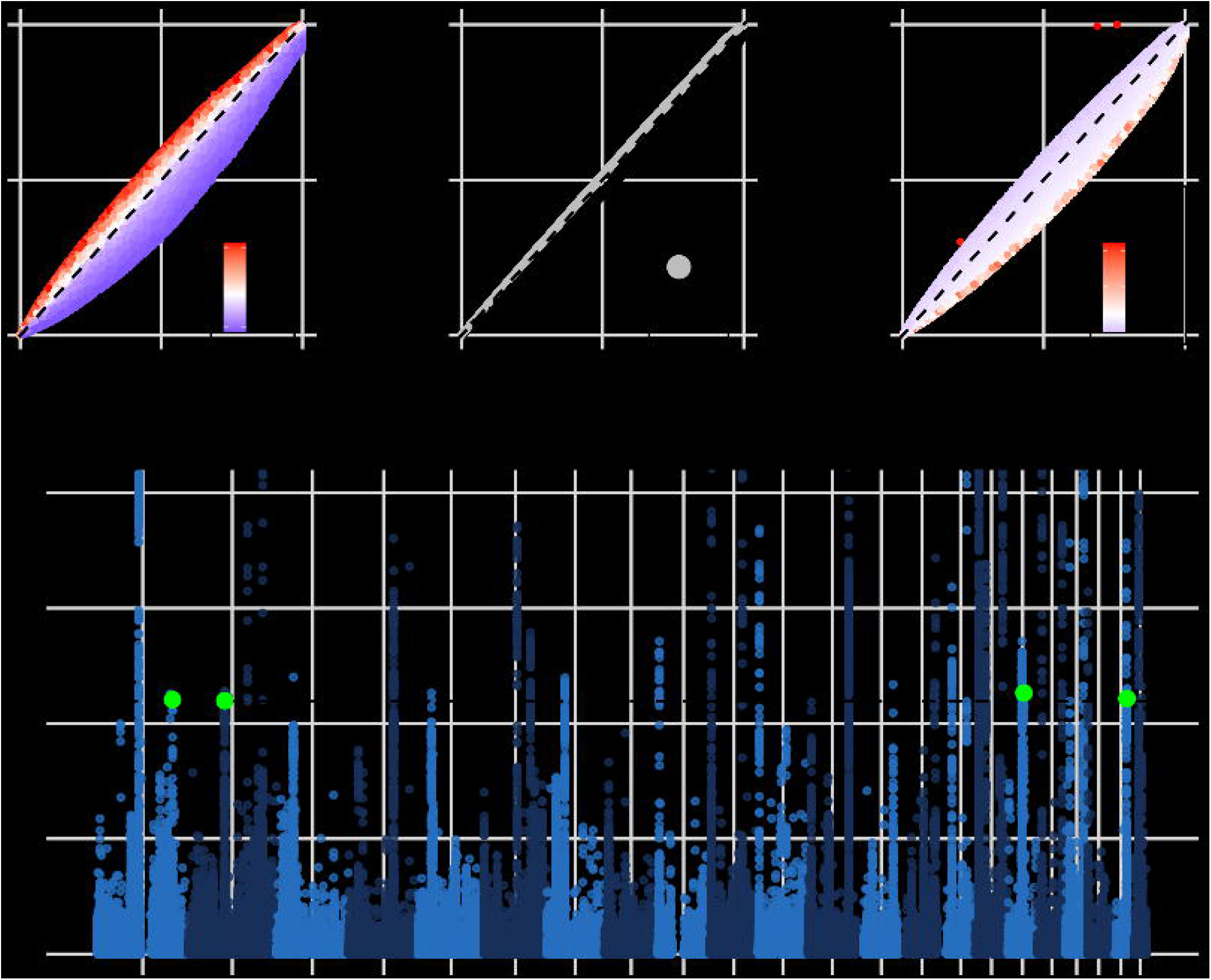

