## Supplemental Material for "fcfdr: an R package to leverage continuous and binary functional genomic data in GWAS"

October 21, 2021

### 1 Supplementary Methods

#### 1.1 Binary cFDR

Let $p_1,...,p_m \in (0,1]$ be a set of $p$-values corresponding to the null hypotheses of no association between the SNP and the trait of interest. Let $q_1,...,q_m \in \{0,1\}$ be a set of binary covariates for the same $m$ SNPs, and denote the null (no association) and alternative (association) hypotheses as $H_0$ and $H_1$ respectively. Assume that $p$ and $q$ are realisations of random variables $P,Q$ satisfying:

$$\begin{aligned} P|H_0 &\sim U(0,1) \\ P &\perp\!\!\!\perp Q|H_0. \end{aligned} \tag{1}$$

We follow the standard methodology introduced by Liley and Wallace (2021) to derive a $v$-value, $v_i$, for each $(p_i,q_i)$ pair. That is, we find the smallest rejection region that each observation $(p_i,q_i)$ is contained in, estimate the distribution of $P,Q$ under the null hypothesis and integrate this distribution over the rejection region to obtain the $v$-value.

Since all $q$ are binary, the support of $P,Q$ is two lines and so the rejection regions are of the form

$$L(p_0,p_1) = (P \leq p_0, Q = 0) \cup (P \leq p_1, Q = 1), \tag{2}$$

where $p_0$ and $p_1$ are unknown.

We wish to find $v$-values such that for all $\alpha$,

$$\begin{aligned} &Pr(v_i < \alpha|H_0) = \alpha \\ &Pr(v_i < \alpha|H_1) \text{ is maximal.} \end{aligned} \tag{3}$$

That is, the $v$-values behave like $p$-value in that they are uniform under the null, but are as small as possible under the alternative hypothesis. Appendix A.1 in Liley and Wallace (2021) (and also Du and Zhang (2014) and Alishahi *et al.* (2016), for example) show that this corresponds to rejection regions formed by the set of points for which $f_0(p,q)/f_1(p,q) < k(\alpha)$, for some $k$, where $f_0(p,q) = f(P=p,Q=q|H_0)$ and $f_1(p,q) = f(P=p,Q=q|H_1)$. That is, $p_0$ and $p_1$ will satisfy the property

$$\frac{f_0(p_0,0)}{f_1(p_0,0)} = \frac{f_0(p_1,1)}{f_1(p_1,1)}. \tag{4}$$

Let

$$f(p,q) = f(P=p,Q=q) = \pi_0 f_0(p,q) + (1-\pi_0)f_1(p,q), \tag{5}$$

where $\pi_0 = Pr(H_0)$. Then equation (4) implies

$$f_0(p_0,0)f_1(p_1,1) = f_0(p_1,1)f_1(p_0,0) \tag{6}$$

$$f_0(p_0,0)\frac{f(p_1,1)-\pi_0 f_0(p_1,1)}{1-\pi_0} = f_0(p_1,1)\frac{f(p_0,0)-\pi_0 f_0(p_0,0)}{1-\pi_0} \tag{7}$$

$$\frac{f(p_1,1)-\pi_0 f_0(p_1,1)}{f_0(p_1,1)} = \frac{f(p_0,0)-\pi_0 f_0(p_0,0)}{f_0(p_0,0)} \tag{8}$$

$$\frac{f(p_1,1)}{f_0(p_1,1)} = \frac{f(p_0,0)}{f_0(p_0,0)}. \tag{9}$$

To solve equation (4) for $p_0$ and $p_1$, we approximate

$$\frac{f_0(p_i,q_i)}{f(p_i,q_i)} = \frac{Pr(P=p_i,Q=q_i|H_0)}{Pr(P=p_i,Q=q_i)} \tag{10}$$

$$\approx \frac{Pr(P\le p_i,Q=q_i|H_0)}{Pr(P\le p_i,Q=q_i)} \tag{11}$$

$$= \frac{Pr(P\le p_i|Q=q_i,H_0)Pr(Q=q_i|H_0)}{Pr(P\le p_i|Q=q_i)Pr(Q=q_i)} \tag{12}$$

$$\approx \frac{p_i \times \widehat{Pr(Q=q_i|H_0)}}{|j:p_j\le p_i,q_j=q_i|/m} \tag{13}$$

where $\widehat{Pr(Q=q_i|H_0)} = \dfrac{|j:q_j=q_i,p_j>1/2|}{|j:p_j>1/2|}$ and $m$ is the total number of observations (i.e. the total number of SNPs). If $q_i=0$ then we set $p_0=p_i$ and use approximation (13) to solve equation (4) for $p_1$. If $q_i=1$, then we set $p_1=p_i$ and solve for $p_0$.

Specifically, if $q_i = 0$ then we set $p_0 = p_i$ and solve the following for $p_1$:

$$\frac{p_i \times \frac{|j : q_j = 0, p_j > 1/2|}{|j : p_j > 1/2|}}{|j : p_j \leq p_i, q_j = 0|/m} = \frac{p_1 \times \frac{|j : q_j = 1, p_j > 1/2|}{|j : p_j > 1/2|}}{|j : p_j \leq p_1, q_j = 1|/m} \tag{14}$$

$$\frac{p_i \times \frac{|j : q_j = 0, p_j > 1/2|}{|j : p_j > 1/2|}}{|j : p_j \leq p_i, q_j = 0| \times \frac{|j : q_j = 1, p_j > 1/2|}{|j : p_j > 1/2|}} = \frac{p_1}{|j : p_j \leq p_1, q_j = 1|}. \tag{15}$$

In practise, we do this using a fold-removal protocol for estimation to ensure that rejection rules are not applied to the same data on which those rules were determined. Specifically, we either leave out each chromosome or each LD block in turn and use the remaining SNPs to estimate the values for the held out SNPs.

Similarly, if $q_i = 1$, then we set $p_1 = p_i$ and solve the following for $p_0$:

$$\frac{p_0 \times \frac{|j : q_j = 0, p_j > 1/2|}{|j : p_j > 1/2|}}{|j : p_j \leq p_0, q_j = 0|/m} = \frac{p_i \times \frac{|j : q_j = 1, p_j > 1/2|}{|j : p_j > 1/2|}}{|j : p_j \leq p_i, q_j = 1|/m} \tag{16}$$

$$\frac{p_0}{|j : p_j \leq p_0, q_j = 0|} = \frac{p_i \times \frac{|j : q_j = 1, p_j > 1/2|}{|j : p_j > 1/2|}}{|j : p_j \leq p_i, q_j = 1| \times \frac{|j : q_j = 0, p_j > 1/2|}{|j : p_j > 1/2|}}. \tag{17}$$

We derive the final $v$-values by integrating the distribution of $P,Q$ under the null hypothesis over the rejection regions:

$$\int_{L(p_0,p_1)} df_0 = Pr((P,Q) \in L(p_0,p_1)|H_0) \tag{18}$$

$$= Pr((P \leq p_0, Q = 0) \cup (P \leq p_1, Q = 1)|H_0) \tag{19}$$

$$\begin{aligned} &= Pr(P \leq p_0, Q = 0|H_0) \\ &\quad + Pr(P \leq p_1, Q = 1|H_0) \end{aligned} \tag{20}$$

$$\begin{aligned} &= Pr(P \leq p_0|Q = 0, H_0)Pr(Q = 0|H_0) \\ &\quad + Pr(P \leq p_1|Q = 1, H_0)Pr(Q = 1|H_0) \end{aligned} \tag{21}$$

$$= p_0 \times (1 - q_0) + p_1 \times q_0 \tag{22}$$

where $q_0 = \widehat{Pr(Q = 1|H_0)}$.

The $v$-value, $v_i$, can be interpreted as the probability that a randomly-chosen $(p,q)$ pair has a more extreme cFDR value than cFDR$(p_i,q_i)$ under $H_0$. That is, a quantity analogous to a $p$-value that can be readily FDR controlled using any FDR controlling procedure that allows for slightly dependent $p$-values, such as the Benjamini-Hochberg (BH) procedure (Benjamini and Hochberg, 1995).

#### 1.2 Simulation analysis

We evaluated the performance of Binary cFDR as implemented in the `fcfdr` R package using a simulation-based analysis.

##### 1.2.1 Simulating GWAS results (p)

Following Hutchinson *et al.* (2021), we first simulated GWAS $p$-values for the arbitrary "principal trait". We collected haplotype data for 3781 individuals from the UK10K project (REL-2012-06-02) (The UK10K Consortium, 2015) at 80,356 SNPs residing on chromosome 22 with $MAF \geq 0.05$ (to match the convention that genetic association studies identify common genetic variation). We split the haplotype data into 24 LD blocks representing approximately independent genomic regions defined by the LD detect method (Berisa and Pickrell, 2016). We then further stratified these so that no more than 1000 SNPs were present in each block, subsequently recording the LD block that each SNP resided in.

We used the `simGWAS` R package (`https://github.com/chr1swallace/simGWAS`) (Fortune and Wallace, 2019) to simulate $Z$-scores for SNPs within each block. The `simGWAS::simulate_z_scores` function requires input for (i) the number of cases and controls (ii) the causal variants (iii) the log ORs at the causal variants and (iv) haplotype frequencies. For our simulation analysis, we selected 5000 cases and 5000 control samples, and within each block we randomly sampled 2, 3 or 4 causal variants with log OR effect sizes simulated from the standard Gaussian prior used in case-control genetic fine-mapping studies, $N(0, 0.2^2)$ (Wellcome Trust Case Control Consortium, 2007). For the haplotype frequency parameter, we supplied a `data.frame` of haplotypes using the UK10K data, with a column of computed frequencies for each haplotype. We collated the $Z$-scores from each region and converted these to $p$-values representing the evidence of association between the SNPs and the arbitrary principal trait.

##### 1.2.2 Simulating auxiliary data (q)

We considered three use-cases of Binary cFDR (simulations A-C) defined by dependence on the principal trait $p$-value ($p_i$) and correlations between realisations of $q$. In simulation A we leveraged binary auxiliary data that was independent of $p_i$: $q_i \sim \text{Bernoulli}(0.05)$. In simulations B and C we leveraged binary auxiliary data that was dependent on $p_i$ by first defining "functional SNPs" as causal variants plus any SNPs within 10,000-bp (to incorporate SNPs residing in the same arbitrary "functional mark"), and "non-functional SNPs" as the remainder. We then sampled $q_i$ from different mixture Gaussian distributions for functional and non-functional SNPs. Specifically, in simulation B we sampled:

$$q_i \sim \begin{cases} \text{Bernoulli}(0.05), \text{ if SNP } i \text{ is non-functional} \\ \text{Bernoulli}(0.4), \text{ if SNP } i \text{ is functional.} \end{cases} \tag{23}$$

Our method will likely be used to leverage functional genomic data iteratively, and so we also evaluated the impact of repeatedly iterating over auxiliary data that captured the same functional mark. Thus, in simulation C we iterated over realisations of $q$ that were highly correlated:

$$q_i \sim \begin{cases} \text{Bernoulli}(0.05), & \text{if SNP } i \text{ is non-functional} \\ \text{Bernoulli}(0.8), & \text{if SNP } i \text{ is functional.} \end{cases} \quad (24)$$

(The auxiliary data is highly correlated in simulation C because in each iteration 80% of the functional SNPs are expected to have an auxiliary data value of 1, and the functional SNPs are the same across iterations in each simulation.)

##### 1.2.3 Implementing Binary cFDR

We used the `fcfdr::binary_cfdr` function to implement Binary cFDR in our simulation analysis. To ensure that rejection rules were not applied to the same data on which those rules were determined, we used a vector of indices of the LD blocks (Berisa and Pickrell, 2016) that each SNP resided on for the `group` parameter. In each simulation for each simulation scenario, we applied Binary cFDR iteratively 5 times to represent leveraging multi-dimensional covariates.

##### 1.2.4 Evaluating sensitivity, specificity and FDR control

To quantify the results from our simulations, we used the BH procedure to derive FDR-adjusted $v$-values from Flexible cFDR, which we call "FDR values" for conciseness (that is, we used the `stats::p.adjust` R function with `method="BH"`). We then calculated proxies for the sensitivity (true positive rate) and the specificity (true negative rate) at an FDR threshold of $\alpha = 5e-06$, which roughly corresponds to the genome-wide significance $p$-value threshold of $5e-08$ (the maximum FDR value amongst SNPs with raw $p$-value $\leq 5e-08$ was $5.4e-06$). We defined a subset of "truly associated SNPs" as any SNPs with $r^2 \geq 0.8$ with any of the causal variants. Similarly, we defined a subset of "truly not-associated SNPs" as any SNPs with $r^2 \leq 0.01$ with all of the causal variants. (Note that there are 3 non-overlapping sets of SNPs: "truly associated", "truly not-associated" and neither of these). We calculated the sensitivity proxy as the proportion of truly associated SNPs that were called significant and the specificity proxy as the proportion of truly not-associated SNPs that were called not significant.

To assess whether the FDR was controlled within a manageable number of simulations, we raised $\alpha$ to 0.05 and calculated the proportion of SNPs called FDR significant which were truly not-associated (that is, $r^2 \leq 0.01$ with all of the simulated causal variants).

#### 1.3 T1D application

##### 1.3.1 T1D GWAS data

We downloaded full harmonised GWAS summary statistics for T1D (Onengut-Gumuscu *et al.*, 2015) from the NHGRI-EBI GWAS Catalog (Buniello *et al.*, 2019) (study GCST005536 accessed on 08/10/21) and used these as the principal trait *p*-values. We then used the LDAK software (`https://dougspeed.com/ldak/`) to obtain LDAK weights for each SNP, and defined our independent SNP set (used to fit the KDE in Flexible cFDR) as the set of SNPs given a non-zero LDAK weight (an LDAK weight of 0 means that its signal is (almost) perfectly captured by neighbouring SNPs).

For the MAF matching step described in Hutchinson *et al.* (2021), we used MAFs estimated from the CEU sub-population samples in the 1000 Genomes Project Phase 3 data set (The 1000 Genomes Project Consortium, 2015). For any SNPs with missing MAF, we randomly sampled a value from the empirical distribution of non-missing MAFs.

To define independent loci for our locus-level results, we first calculated LD between each pair of SNPs using haplotype data from the 503 individuals of European ancestry in the 1000 Genomes Project Phase 3 data set (The 1000 Genomes Project Consortium, 2015). We then used PLINK's LD-clumping algorithm with a 5-Mb window and an $r^2$ threshold of 0.01. This conservative clumping approach sorts SNPs into ascending order of *p*-value and then moves down the list, sequentially removing SNPs within a 5-Mb window and with $r^2 > 0.01$. The SNP with the smallest *p*-value in the data set in each LD clump was called the "lead variant".

##### 1.3.2 Validation GWAS data set

We downloaded full harmonised GWAS summary statistics for T1D (Robertson *et al.*, 2021) from the NHGRI-EBI GWAS Catalog (Buniello *et al.*, 2019) (study GCST90013445 accessed on 08/10/21) and used this as our validation GWAS data set. The samples in the discovery GWAS data set (Onengut-Gumuscu *et al.*, 2015) were a subset of those in the validation data set, and so we said that a discovery validated if it's corresponding $p$-value was smaller in Robertson *et al.* (2021) than Onengut-Gumuscu *et al.* (2015).

##### 1.3.3 Auxiliary data

We downloaded full harmonised GWAS summary statistics for rheumatoid arthritis (RA) (Eyre *et al.*, 2012) from the NHGRI-EBI GWAS Catalog (Buniello *et al.*, 2019) (study GCST005569 accessed on 08/10/21). We mapped each SNP in the T1D GWAS data set to its corresponding $p$-value for RA using genomic coordinates and rsIDs. We removed 6044 SNPs from the analysis which did not have a corresponding $p$-value for RA.

We downloaded SNP-level annotations for all 1000 Genomes SNPs from the baseline-LD model (version 2.2) described in Gazal *et al.* (2017). We extracted values for the binary annotation "DGF_ENCODE" which quantifies sites of transcription factor occupancy. Briefly, this annotation is derived from merging all DNase I digital genomic footprinting (DGF) regions from the narrow-peak classifications across 57 cell types (ENCODE Project Consortium, 2012; Gusev *et al.*, 2014). DGF regions (corresponding to DGF annotation values of 1) are expected to precisely map sites where regulatory factors bind to the genome (Neph *et al.*, 2012). We matched each SNP in the T1D GWAS data set to its binary DGF annotation using genomic coordinates. We removed 2811 SNPs from the analysis that did not have a corresponding DGF annotation value.

We downloaded consolidated fold-enrichment ratios of H3K27ac ChIP-seq counts relative to expected background counts from NIH Roadmap Epigenomics Mapping Consortium (Bernstein *et al.*, 2010) in nine primary tissues and cells relevant for T1D (CD3, CD4+ CD25int CD127+ Tmem, CD4+ CD25+ CD127- Treg, CD4+ CD25- Th, CD4+ CD25- CD45RA+, CD4 memory, CD4 naive, CD8 memory, CD8 naive). Specifically, we downloaded the `bigWig` files from `https://egg2.wustl.edu/roadmap/data/byFileType/signal/consolidated/macs2signal/foldChange/`, converted these to `wig` files and then to `bed` files, and then mapped each SNP in the T1D GWAS data set to its corresponding genomic region in the `bed` files and recorded the H3K27ac fold change values in each cell type using the `bedtools intersect` utility. For SNPs on the boundary of a genomic region (and therefore mapping to two regions) we randomly selected one of the regions. We observed that the fold change values across T1D-relevant cell types were highly correlated ($r > 0.65$) (Supplementary Fig. 1) and therefore averaged values across cell types to avoid iterating over highly correlated auxiliary data that is likely capturing the same functional mark. We transformed the averaged fold change values ($q := log(q+1)$) to deal with long tails.

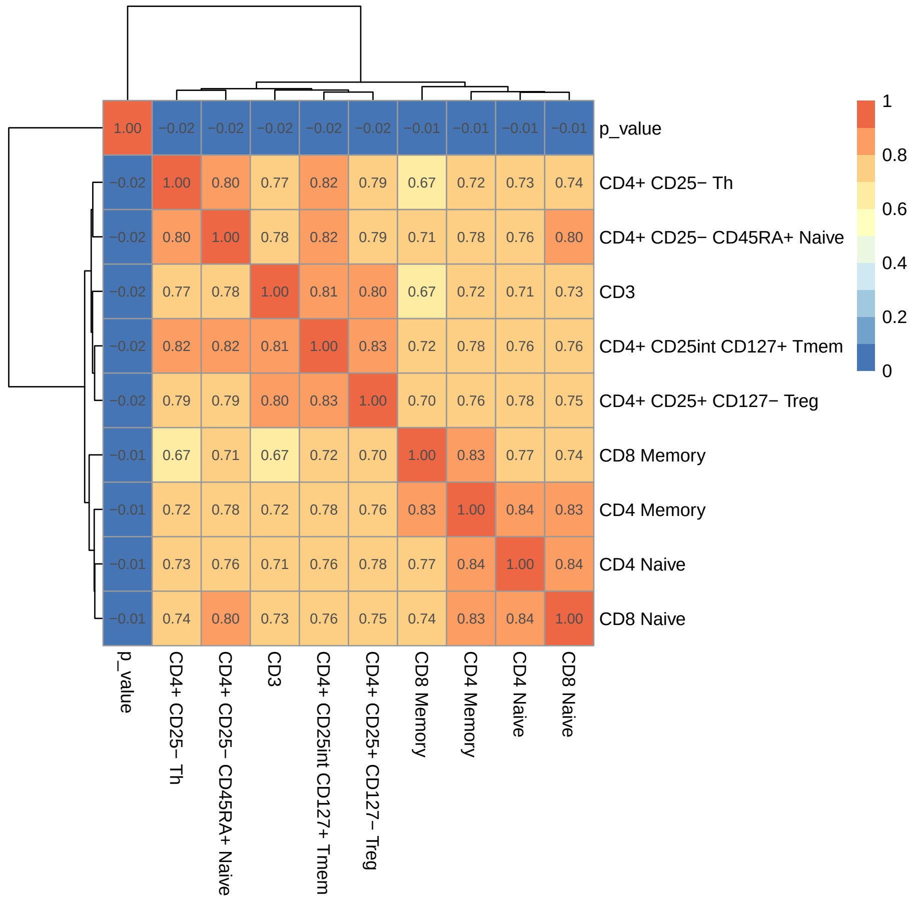

Supplementary Fig. 1: Heatmap of the Pearson correlation coefficients between H3K27ac fold change values amongst T1D-relevant cell types downloaded from NIH Roadmap Epigenomics Mapping Consortium (Bernstein *et al.*, 2010) and (log transformed) T1D $p$-values from Onengut-Gumuscu *et al.* (2015). Figure generated using the `pheatmap` R package (`https://cran.r-project.org/web/packages/pheatmap/index.html`).

##### 1.3.4 Implementation

We used the `fcfdr::flexible_cfdr` and `fcfdr::binary_cfdr` functions to leverage the auxiliary data with T1D GWAS $p$-values iteratively. We used the chromosome for which each SNP resided for the `group` parameter in `fcfdr::binary_cfdr`, and we used the estimated MAF values for the optional `maf` parameter in the `fcfdr::flexible_cfdr` function. We used the `stats::p.adjust` function with `method="BH"` to derive FDR values from the $v$-values (after the 3 iterations) and used these as the output of interest. We used an FDR threshold of

$FDR \leq 3.305365e-06$ to call significant SNPs, which corresponded to the genome-wide significance threshold $p \leq 5e-08$ (it was the maximum FDR value amongst SNPs with raw $p$-values $\leq 5e-08$ in the discovery GWAS data set). The full data and code to replicate the analysis are available from `https://annahutch.github.io/fcfdr/articles/t1d_app.html`.

### 2 Supplementary Results

#### 2.1 Results from simulation analysis

We expect that leveraging irrelevant data should not change our conclusions about a study. Simulation A showed that the sensitivity and specificity remained stable across iterations and that the FDR was controlled at a pre-defined level when using Binary cFDR to leverage independent binary auxiliary data with arbitrary GWAS $p$-values (Supplementary Fig. 2A). In contrast, when leveraging relevant data we hope that the sensitivity improves whilst the specificity remains high. This is what we observed for Binary cFDR in simulation B (Supplementary Fig. 2B).

It is known that cFDR should not be used to iterate over highly correlated auxiliary data that is capturing the same functional mark, as SNPs with a modest $p$ but extreme $q$ will incorrectly attain greater significance with each iteration (for a more detailed explanation see Hutchinson *et al.* (2021)). Simulation C involved iterating over highly correlated auxiliary data values (mean Pearson correlation coefficient was 0.3) that capture the same "functional mark" (80% of functional SNPs were expected to have an auxiliary data value of 1 in each iteration). The lack of FDR control in simulation C (Supplementary Fig. 2C) serves as a salutary reminder that care should be taken not to repeatedly iterate over functional data that is capturing the same genomic feature.

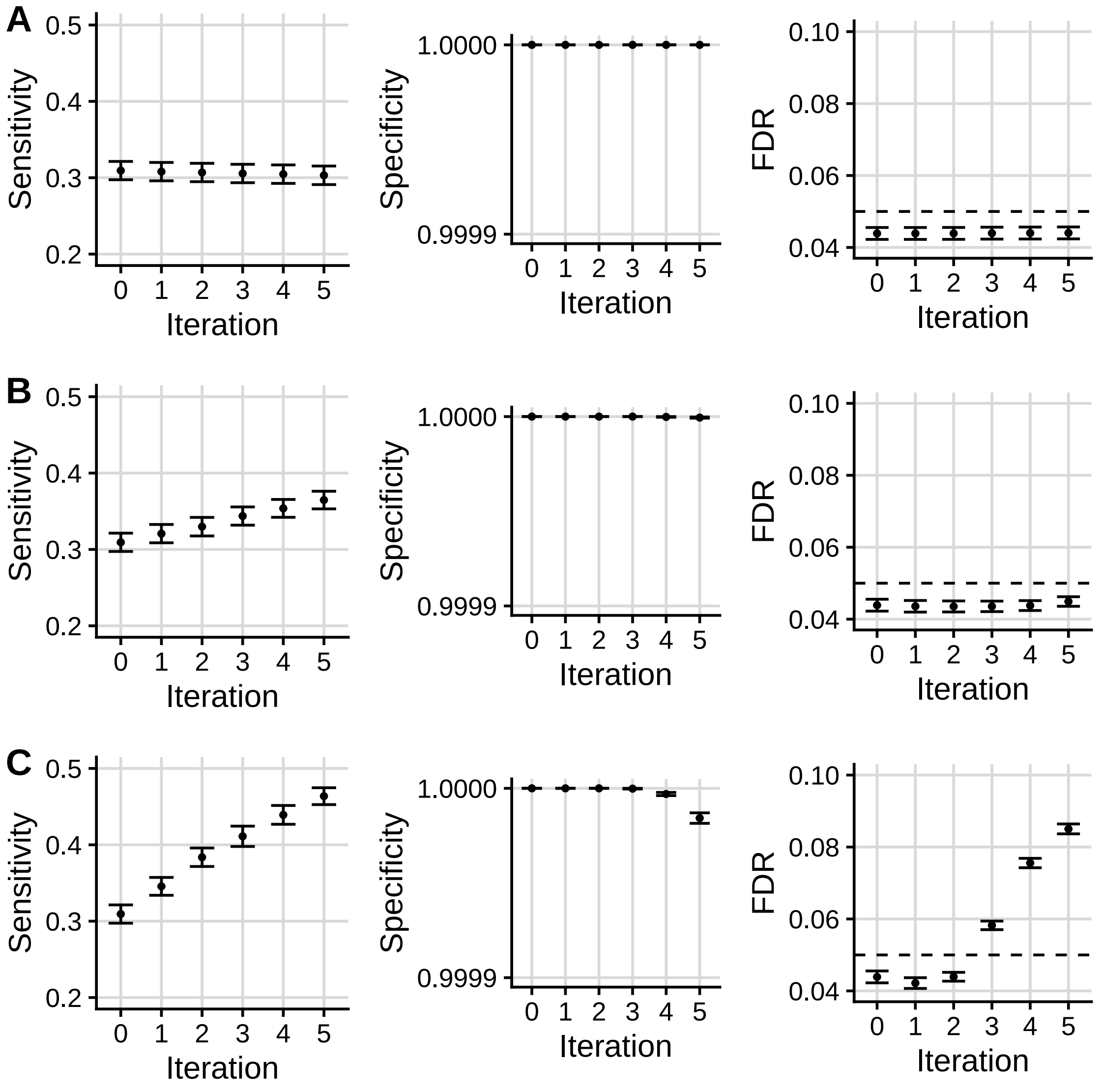

Supplementary Fig. 2: Mean +/- standard error for the sensitivity, specificity and FDR of FDR values from Binary cFDR when iterating over independent (A; "simulation A") and dependent (B; "simulation B" and C; "simulation C") binary auxiliary data. Iteration 0 corresponds to the original FDR values. Our sensitivity proxy is calculated as the proportion of SNPs with $r^2 \geq 0.8$ with a causal variant ("truly associated"), that were detected with a FDR value less than $5e-06$. Our specificity proxy is calculated as the proportion of SNPs with $r^2 \leq 0.01$ with all the causal variants ("truly not-associated"), that were not detected with a FDR value less than $5e-06$. Our FDR proxy is calculated as the proportion of SNPs that were detected with a FDR value less than 0.05, that had $r^2 \leq 0.01$ with all the causal variants ("truly not-associated") (we raised $\alpha$ to 0.05 in order to assess FDR control within a manageable number of simulations). Results were averaged across 100 simulations.

##### 2.2 Supplementary results from T1D application

We evaluated the relationship between the "principal $p$-values" ($p$) and the auxiliary data ($q$) in each iteration. In iteration 1 the Pearson correlation coefficient between $p$ (T1D GWAS $p$-values) and $q$ (RA GWAS $p$-values) was 0.092 (Supplementary Fig. 3A). In iteration 2 the Pearson correlation coefficient between $p$ ($v$-values from iteration 1) and $q$ (binary DGF value) was -0.022. In iteration 3 the Pearson correlation coefficient between $p$ ($v$-values from iteration 2) and $q$ (log transformed average H3K27ac counts) was -0.083 (Supplementary Fig. 3B).

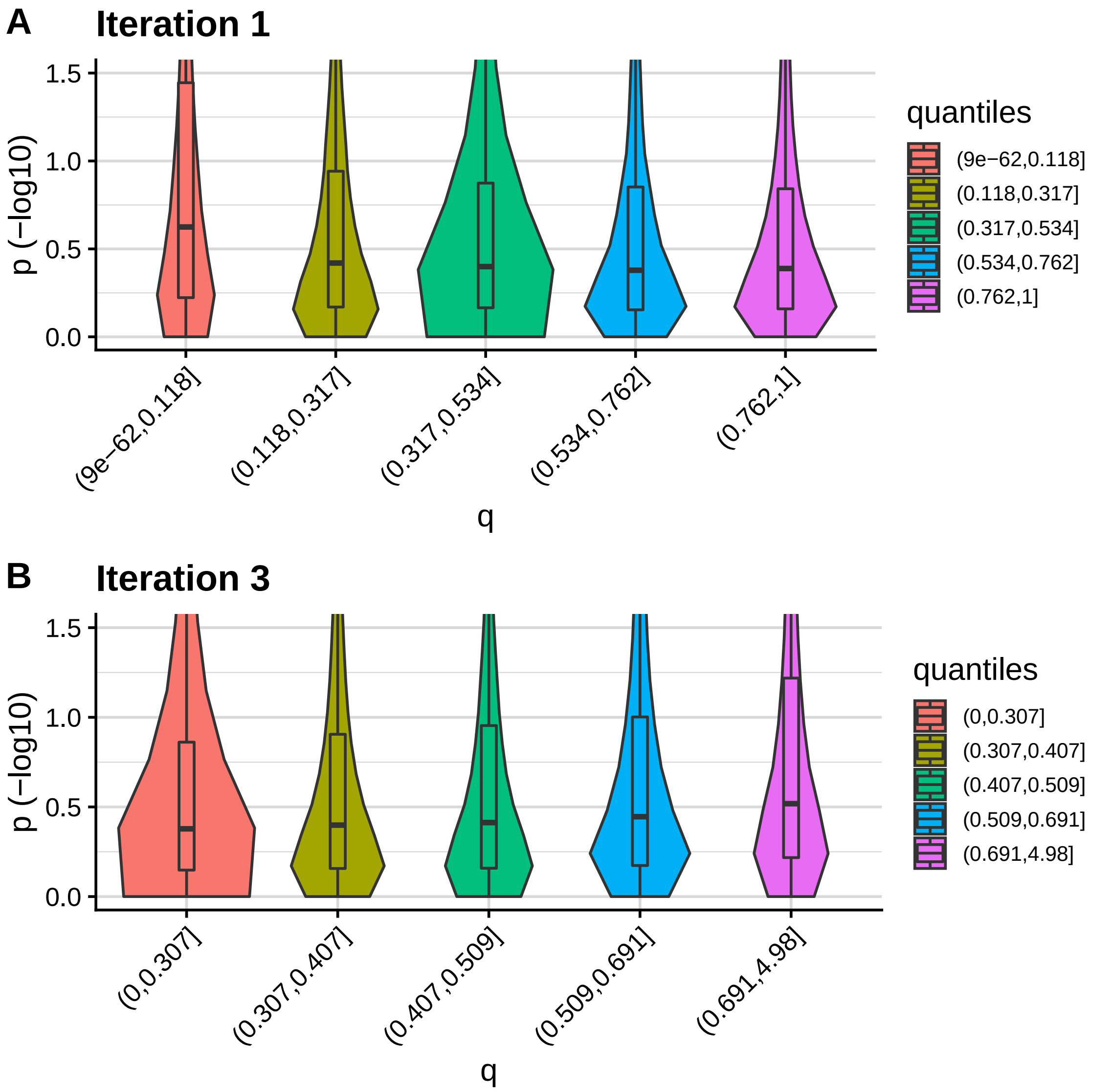

Supplementary Fig. 3: Violin plots showing the relationship between $p$ and $q$ in iterations 1 and 3 of the cFDR framework in the T1D application. Figure generated using the `fcfdr::corr_plot` function with default parameter values (`https://annahutch.github.io/fcfdr/reference/corr_plot.html`).
